# Enhanced BCMA Antigen Density Increases Trogocytosis and Attenuates CAR T cell Function

**DOI:** 10.64898/2026.07.15.738803

**Authors:** Pinar Ataca Atilla, Sylvain Simon, Erden Atilla, David Coffey, Andrew J Cowan, Grace Bugos, Thomas J Diefenbach, Janice Huang, Efe Karaca, Zhiqun Zhou, Melissa L Comstock, Geoffrey R Hill, Stanley R Riddell, Damian J Green

## Abstract

**Background:** Chimeric Antigen Receptor (CAR) T cell therapy targeting B-cell maturation antigen (BCMA) has demonstrated impressive clinical efficacy in relapsed/refractory multiple myeloma (MM). Nonetheless, disease relapse limits durable response for most patients. Trogocytosis of target antigen by effector cells has emerged as a potential contributor to reduced surface antigen density, CAR T cell dysfunction, and fratricide. Although γ-secretase inhibitors (GSI) significantly increase cell surface BCMA density and decrease soluble BCMA (sBCMA), their effects on BCMA trogocytosis and the resulting impact on CAR T-cell function remain incompletely understood.

**Methods:** We investigated the effects of GSI on BCMA-directed CAR T cell function and trogocytosis using in vitro co-culture systems with MM cell lines across a spectrum of BCMA expression. We validated findings using confocal microscopy and cytotoxicity assays. Trogocytosis and fratricide were assessed in time-resolved functional studies. Phenotypic and functional differences between trogocytosis-positive (CAR T Trogo+) and trogocytosis-negative (CAR T) cells were evaluated using multiparametric flow cytometry, proteomic profiling, single-cell RNA sequencing (scRNA-seq), T-cell receptor (TCR) sequencing, and in vitro rechallenge assays. We also interrogated clinical samples from two Phase I trials (NCT03338972 and NCT03502577) which employed the identical CAR T cell construct with or without GSI respectively, to evaluate the relationship between trogocytosis, CAR T cell persistence, and treatment outcome.

**Results:** GSI driven increases in BCMA density on MM cell lines enhanced CAR T cell cytotoxicity but concomitantly increased trogocytosis, particularly in high-antigen-density cell lines-(H929+GSI *vs* H929; 30 min (P<0.0001), 1 h (P<0.0001), 2 h (P<0.0001), 6 h (P<0.0001), and 24 h (P<0.0001) and in CD4⁺ CAR T cells (K562mCherry+GSI, CD4 *vs* CD8 CAR T cells;10 min (P=0.01), 2 h (P=0.01), and 6 h (P=0.004). Following BCMA acquisition, CAR T cells (CAR T Trogo⁺) exhibited reduced proliferative capacity, diminished cytotoxic function (CAR T Trogo+ vs CAR T; (P=0.01), and an increase in markers of exhaustion/activation (CD4+ CAR T Trogo+ *vs* CD4 CAR T and CD8+ CAR T Trogo+ *vs* CAR T; PD-1+LAG-3+, PD-1+TIM-3+, and TOX+TIM-3 co-expression, (P=0.007, P<0.0001, P=0.006 and P=0.004, P=0.03, P=0.006). In fratricide assays, CAR T Trogo⁺ cells were susceptible to killing by naïve CAR T cells. Single cell RNA-seq supports the phenotypic findings revealing transcriptional features of heightened activation and accelerated exhaustion in CAR T Trogo+ cells. Clinical phase I trial data confirm BCMA trogocytosis in patient samples.

**Conclusions:** Our findings highlight the paradoxical effects of increased BCMA density on BCMA CAR T cell therapy: enhancement of initial tumor targeting and promotion of trogocytosis-associated dysfunction. Trogocytosis may contribute to antigen modulation, CAR T cell exhaustion, and fratricide, potentially muting the therapeutic benefits of enhanced antigen density. To optimize GSI and mitigate trogocytosis-associated resistance mechanism, future clinical trial designs should incorporate early time-point sampling, a sample size providing sufficient statistical power to determine an impact on CAR T cell persistence and treatment response, and mechanistic assessments.

## Introduction

Chimeric Antigen Receptor (CAR) T cell therapies provide highly effective, targeted treatment options that have achieved remarkable clinical responses in otherwise treatment-resistant B cell malignancies. Targeting B-cell maturation antigen (BCMA) with CAR T cells in multiple myeloma (MM) has emerged a promising strategy reaching 73-100% overall response rate (ORR) in patients after ≥ three prior lines of therapy (1–3). While long-term follow-up of these heavily pretreated cohorts has demonstrated durable remissions in a subset of patients, disease relapse remains the predominant cause of treatment failure following BCMA CAR T cell therapy (6). More recently, both ide-cel and cilta-cel have been approved in earlier lines of therapy (4,5). Whether treatment earlier in the disease course will improve the long-term durability of response remains to be determined.

Tumor antigen escape represents a key mechanism for resistance to CAR-T therapy (7,8). With the application of whole genome sequencing (WGS) and single cell RNA-sequencing (scRNA-seq) the molecular mechanisms underlying antigen escape following BCMA CAR T cell therapy have been better characterized. Although complete loss of BCMA expression is relatively rare in relapsed patients following BCMA CAR-T cell therapy, pre-existing subclones harboring monoallelic 16p deletion, frequently observed in newly diagnosed high-risk MM, may predispose to biallelic inactivation of *TNFRSF17* (encoding BCMA). This loss can occur through clonal selection or antigenic drift (9–11). In contrast to CD19, BCMA-targeted therapies are affected by the dynamic interplay between soluble BCMA and its membrane-bound form on the tumor cell surface. The γ secretase (GS) complex mediates proteolytic cleavage of the BCMA receptor, limiting BCMA surface accumulation and increasing levels of soluble BCMA (sBCMA). Soluble BCMA impairs humoral immune responses by antagonizing BAFF-mediated B cell survival signaling (12–14). The successful blockage of BCMA cleavage by γ secretase inhibition enhances CAR T cell efficacy preclinically (15) supporting a first-in-human Phase I trial with BCMA CAR T cell therapy in combination with oral GSI, crenigacestat (NCT03502577), in heavily pre-treated high-risk patients. After 3 oral doses of GSI, BCMA antibody binding capacity increased by a median of 12.2-fold, accompanied by a reduction in sBCMA levels. At a median follow-up of 36 months (95% Cl, 26 to not reached), the PFS for all patients was 11 months (95% Cl, 5.4 to not reached) with superior outcomes amongst BCMA naïve patients (16).

Despite the heavily pre-treated (median ten prior regimens) and high-risk population, the degree of enhanced target density was not proportional to response duration. This prompted tour studies investigating alternative mechanisms underlying relapses following combined GSI and BCMA CAR T cell therapy. Higher levels of trogocytosis have contributed to relapse in various CAR T/NK models (17–19). Fundamentally, trogocytosis involves rapid extraction of membrane fragments including surface antigens during direct cell-to-cell contact (20). In this study, we aimed to investigate the impact of GSI on BCMA CAR T cell trogocytosis. We characterized trogocytosis, fratricide, CAR T cell phenotype and function, and the molecular features of trogocytosis-positive CAR T cells using functional assays, multiparametric flow cytometry, proteomic profiling and confocal microscopy. Finally, we analyzed data from two Phase I BCMA CAR T cell clinical trials (NCT03338972 and NCT03502577) to delineate the role of trogocytosis and its modulation by GSI.

## Methods

### Cell lines

Lenti-X cells (Clontech) were cultured as described (21). H929, U266 and wild type K562 were purchased from ATCC and maintained in RPMI 1640 (Gibco-BRL, San Francisco, CA) supplemented with 100 U/mL penicillin/streptomycin (Gibco-BRL, San Francisco, CA) and 10% to 20% fetal bovine serum (Hyclone, Waltham, MA). MOLP-8 was obtained from Deutsche Sammlung von Mikroorganismen and Zellkulturen GmbH, Germany and cultured in RPMI 1640 supplemented with 100 U/mL penicillin/streptomycin and 20% fetal bovine serum. H929, U266 and MOLP-8 lines were transduced with a gammaretroviral vector encoding enhanced GFP-FFluc fusion protein as previously described (22). Wild type K562 cell line transduced with a lentiviral vector encoding human BCMA-mCherry and sorted regarding to BCMA/mCherry expression. Cells were incubated with GSI, LY3039478-crenigacestat (Eli Lilly) 0.1 μM for 24h as previously described (15).

### Lentiviral vectors, transduction of T cells and functional assays

Healthy donor T cells (Bloodworks Network, Seattle, USA) were activated, transduced with fully human lentiviral BCMA-specific 4-1BB/CD3ζ CAR containing truncated epidermal growth factor receptor (EGFRt), enriched and expanded as described previously (15). After 10 days of expansion, CAR T cells were sorted (BD FACSymphony S6) following staining with anti-EGFR (AY13)(Biolegend, USA). Surface CAR expression was detected by biotinylated recombinant human BCMA protein (ACROBiosystems, USA)+ Streptavidin-APC. Transduced or non-transduced T cells were co-cultured with each of the tumor cell lines described at an E:T ratio in 96 well plates in the absence of exogenous cytokines for 24 hours. We identified tumor cells by GFP and/or mCherry expression. We stained dead cells by measuring the population positive for 7-aminoactinomycin D (7-AAD) (Thermo Fisher Scientific, Waltham, MA) and counted them using CountBright Beads (Thermo Fisher Scientific, Waltham MA). For Incucyte (Sartorius, Gottingen, Germany) assay, tumor cells (K562BCMA-mCherry+/−GSI) and transduced or non-transduced T cells were co-cultured in poly-l-lysine coated plates (Corning, USA) and scanned every 6 hours. Normalized red object count was plotted.

To identify trogocytosis, samples were analyzed by flow cytometry (BD Symphony 5) after combining MM cell line and CAR T cell co-culture for 24hours (h), 6h, 2h, 1h, 30 minutes (min), 10 min (effector to target ratio=1:1). For trogocytosis inhibition assays, CAR T cells were pre-treated with 1 μM latrunculin A (Sigma-Aldrich) at 37 °C for 15 min before co-incubation with target cells.

For fratricide assays, CAR T cells with trogocytosis were sorted following 6h of co-culture with cell lines and re-plated with fresh CAR T cells and were analyzed by flow cytometry after 24h (effector to target ratio=1:1). Soluble BCMA (sBCMA) ELISA was performed according to the manufacturer’s recommendations (R&D Systems).

### Immunophenotyping

T cells were stained with mAbs specific for CD4 (RPA-T4), CD8 (RPA-T8), CD3 (UCHT1) and EGFR (AY13), CD197 (G043H7), CD223 (C9B7W) (Biolegend, USA), CD45RA(HI100), CD279(EH12), CD366 (344823) (BD Biosciences, USA). Cell lines were stained with mAbs specific for BCMA (19F2), CD138 (MI15), CD45 (HI30), CD19 (HIB19), or corresponding isotype controls. Mean fluorescence intensity (MFI) of BCMA on primary MM samples was corrected for isotype staining for GSI-treated or not conditions. For intracellular staining, cells were fixed and permeabilized using the FoxP3 staining kit (ThermoFisher) and stained with mAbs specific for BCMA (19F2). Flow cytometry was performed on a BD FACS Symphony 5 or BD FACS Celesta and data were analyzed with FlowJo (Treestar).

### Confocal Microscopy

CAR T cells and K562BCMA-mCherry+GSI cells were seeded at 1:1 ratio onto poly-L-lysine-coated glass surface chamber slides. Cells were then co-cultured at 37 °C for 1 h. Cells were fixed by adding of 4% paraformaldehyde into the culture medium (final concentration 1%) and incubating for 15 min. After fixation cells were washed twice with PBS. Alexa Flour 555 Anti-EGFR (EP38Y) or Alexa Flour 627 was used for CAR T cell staining. Cells were kept in the dark at −20 °C. Plates were imaged on a Molecular Devices (San Jose, CA) ImageXpress Micro high-content imaging system equipped with a Nikon Plan Fluor ELWD 20x/0.45 objective and an incubation system for maintaining cell viability. Sixteen fields were acquired per well in phase contrast, green fluorescence (CAR-T cells), and red fluorescence (K562BCMA-mCherry+GSI). Images were taken every 15 minutes for 6 hours. Regions of interest highlighting CAR-T interactions with their target cells were chosen manually, and a custom Fiji/ImageJ script was used to assemble single-channel and overlay images, with CAR-T cells outlined.

### Single-cell RNA and TCR sequencing

**a)** **Single-Cell Library Preparation:** Single-cell RNA and TCR libraries were generated using the Parse Biosciences Evercode Mega TCR v3 kit, which utilizes a split-pool combinatorial barcoding strategy. Sorted cells were fixed and permeabilized, then subjected to three rounds of split-pool barcoding to append unique combinatorial barcodes to the cDNA of individual cells. For TCR profiling, targeted enrichment of the V(D)J regions of the T cell receptor alpha and beta chains was performed post-cDNA synthesis. The resulting libraries were sequenced on an Illumina platform.
**b)** **Primary Data Processing and Quality Control:** Raw sequencing reads were processed using the Parse Biosciences TrailMaker™ pipeline (Parse Biosciences, Seattle, USA; available at https://app.trailmaker.parsebiosciences.com; analysis completed on DAT) (22). This included barcode demultiplexing, read alignment to the GRCh38 human reference genome supplemented with the CAR transgenic sequence, and unique molecular identifier (UMI) deduplication. The pipeline generated a filtered gene-barcode count matrix and a matched TCR contig file. Initial quality control in R (v4.4.2) using Seurat (v5.0.1) involved the removal of low-quality cells (23). Cells were retained if they exhibited fewer than 20% mitochondrial-derived transcripts, ensuring the exclusion of apoptotic cells while preserving the high-quality T cell population.
**c)** **Automated Cell Type Annotation and Exhaustion Mapping:** Broad cell types were annotated using CellTypist, utilizing the ‘Immune_All_High’ model (24). This model, trained on high-resolution immune cell atlases, allows for the precise identification of T cells. To achieve mapping of T cell exhaustion and differentiation states, we employed scAtlasVAE (25). This variational autoencoder (VAE) framework utilizes transferring learning to project query cells into a latent space defined by a comprehensive pan-disease CD8+ T cell atlas.
**d)** **Integration and Dimensionality Reduction:** To account for technical batch effects across the six experimental samples, we utilized the Harmony integration algorithm (26). Following integration, we performed uniform manifold approximation and projection (UMAP) on the first 30 Harmony-corrected components for visualization. Due to variability in sequencing depth, samples were downsampled to 1,000 high-quality T cells post-filtering, with gene counts normalized to a median of 300 genes per cell.
**f)** **TCR Clonal Analysis:** T cell receptor diversity was quantified using the Shannon entropy index, which accounts for both the number of unique clonotypes (richness) and the evenness of their distribution.

### Patient Samples

Patient samples from Phase I clinical trial of FCARH143, a fully human BCMA CAR T cell therapy (NCT03338972) (2) and Phase I clinical trial combining crenigacestat with FCARH143 (NCT03502577) manufactured by the Fred Hutchinson Therapeutic Products Program (Seattle, WA) were included in this study (16). We analyzed and normalized mean fluorescence intensity (MFI) of trogocytosed BCMA in CAR T cells by flow cytometry using day 14 and day 28 peripheral blood as well as bone marrow samples. CAR persistence is determined by qPCR and Soluble BCMA (sBCMA) levels were analyzed by ELISA.

### Statistics

For continuous variables, normality was assessed prior to statistical analysis. Comparisons between two groups were conducted using an unpaired two-tailed t-test when data were normally distributed; otherwise, the Mann–Whitney U test was employed. For multiple group comparisons, repeated measures one-way of two-way analysis of variance (ANOVA) followed by Tukey’s post hoc test was used. Statistical comparisons of diversity metrics and subtype abundances between conditions were performed using Student’s t-tests and Fisher’s exact tests, respectively. Statistical analyses were performed using GraphPad Prism (version 8.1.2 or later) or R (version 4.6) or Python (version 3.14.5). P-values <0.05 were considered statistically significant, and significance levels are indicated in the corresponding figures and figure legends.

## Results

We evaluated three multiple myeloma (MM) cell lines characterized by varying BCMA surface density: H929 (high BCMA; mean fluorescence intensity [MFI]: 13,924; antibody binding capacity [ABC]: 26,050), U266 (medium BCMA; MFI: 4,955; ABC:7,210), and MOLP8 (low BCMA; MFI: 2,400; ABC:1,975). Following 24 hours of incubation with the GSI, BCMA surface expression increased approximately 5-fold in H929, 10-fold in U266, and approximately 7-fold in MOLP8 cells (Figure 1A). BCMA CAR successfully transduced on T cells (Suppl Fig 1A) demonstrated significant anti-tumor activity when co-cultured +/− GSI H929 (P<0.0001; P<0.0001), U266 (P<0.0001; P<0.0001) and MOLP8 (P=0.0002; P=0.003) (Suppl Fig 1B, 1C, 1D). Assessment of BCMA acquisition on CAR T cells during co-culture at 10 minutes (min), 30 min, 1 hour (h), 2 h, 6 h, 24 h, demonstrated a time-dependent increase in CAR T cell BCMA MFI across all cell lines, consistent with trogocytosis. GSI treatment, significantly increased trogocytosis in all cell lines evaluated, most strikingly in co-cultures with the H929 cell line, where increased BCMA MFI on CAR T cells is observed at 30 min (P<0.0001), 1 h (P<0.0001), 2 h (P<0.0001), 6 h (P<0.0001), and 24 h (P<0.0001) (Figure 1B). In U266, BCMA MFI was significantly elevated at 30 min (P<0.0001), 1 h (P<0.0001), 2 h (P<0.0001), 6 h (P<0.0001), and 24 h (P=0.002). In MOLP8, significant increases were noted at 30 min (P=0.03), 1 h (P=0.03), and 2 h (P=0.002). Moreover, GSI induced trogocytosis resulted in significantly reduced BCMA surface density on tumor cells, particularly in the H929 cell line (P<0.0001) (Figure 1C). Soluble BCMA (sBCMA) levels concomitantly decreased in H929 cultures treated with GSI (P<0.0001). Concurrently, CAR molecules were also transferred to H929 cell line, and this transfer was higher with GSI (P<0.0001) (Suppl Fig 2A). The loss of surface CAR molecules on CAR T cells after co-culture was significantly greater with GSI at 6h following co-culture in comparison to the same experimental conditions without GSI, potentially reflecting enhanced CAR internalization and/or increased trogocytic transfer of CAR molecules (Suppl Fig 2B). By contrast, the expression of non-cognate antigen CD138 remained unchanged in H929 cells +/− GSI (Suppl Fig 2C).

**Figure 1A.**
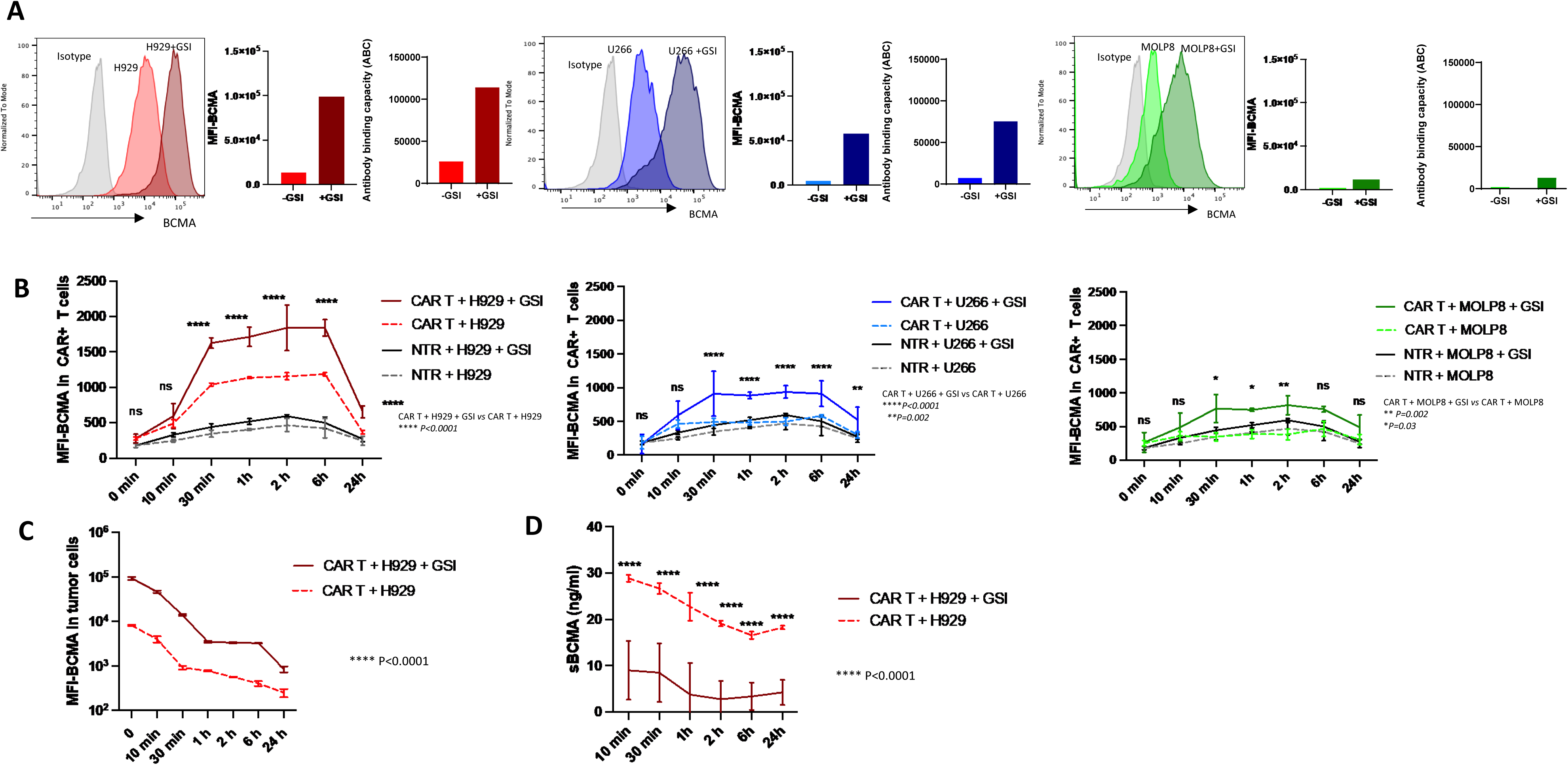
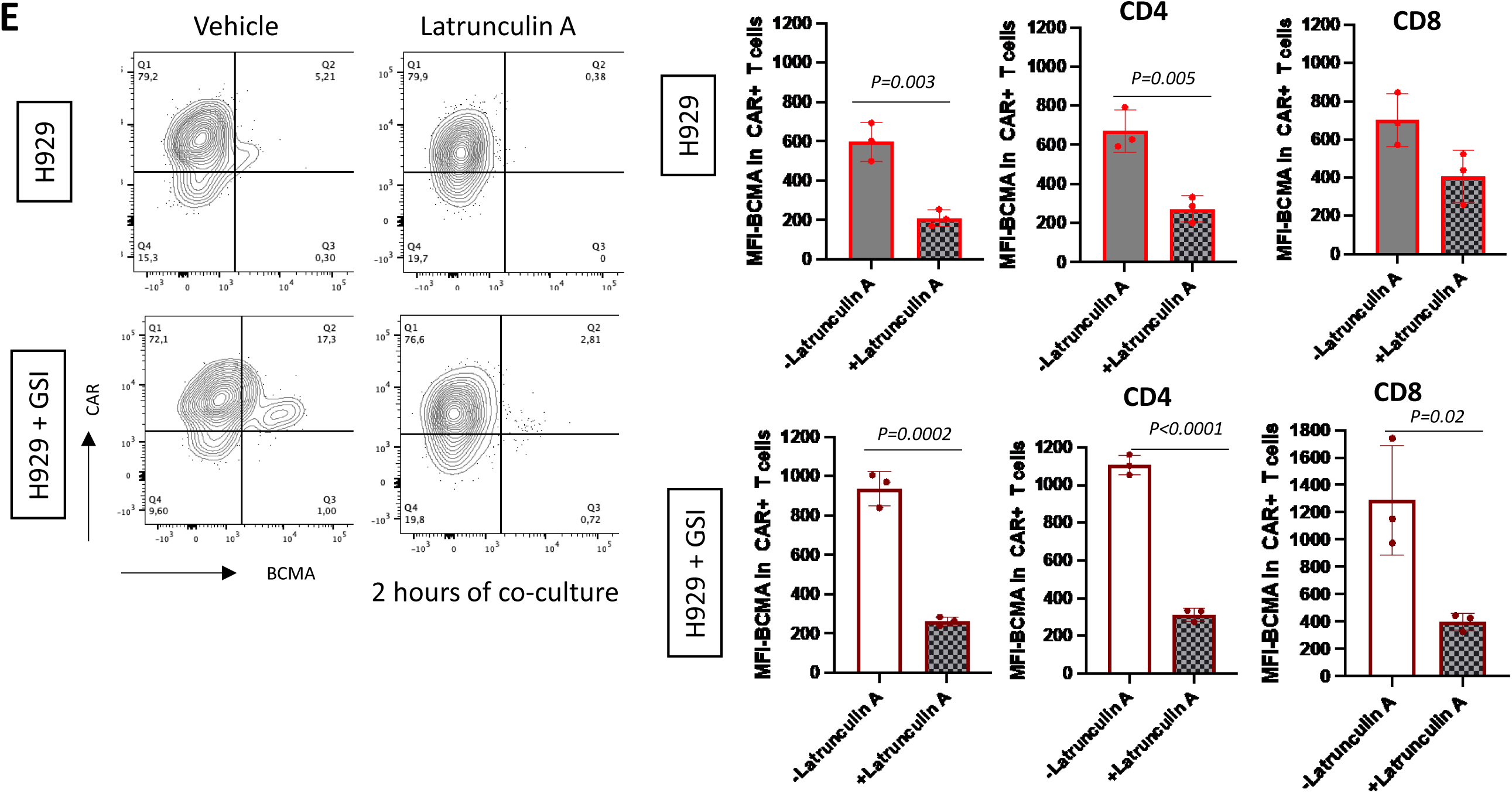
The BCMA expression and MFI-BCMA and ABC-BCMA of cell lines H929 +/−GSI; U266 +/− GSI; MOLP8 +/− GSI **1B.** The BCMA-MFI in CAR+ T and non-transduced T cells in co-cultures with H929 +/−GSI; U266+/− GSI; MOLP8+/− GSI (E:T=1:1) through 24 hours. Two-way ANOVA Tukey’s **1C.** The MFI-BCMA of tumor cells in co-cultures with H929 +/−GSI **1D.** Soluble BCMA (sBCMA) levels in co-cultures with H929+/− (E: Effector, GSI: Gamma secretase inhibitor, T: Target, ns: not significant, NTR:Non-transduced) **Figure 1E**. Trogocytosis is blocked by latrunculin A at 2h of co-culture (E:T=1:1) both in all, CD4 and CD8 CAR T cells H929 +/− GSI

Consistent with the actin-dependent nature of trogocytosis, treatment with latrunculin A, an actin polymerization inhibitor that disrupts F-actin assembly and has been widely used to block trogocytosis, significantly reduced trogocytosis at 2 hours in H929 co-cultures (P=0.002), with the greatest inhibition observed under GSI-treated conditions (P=0.0002), where BCMA MFI on CD4+ CAR T cells was significantly decreased following co-culture with H929 (P=0.005) and further reduced in the presence of GSI (P<0.0001) (17).The reduction in BCMA MFI on CD8+ CAR T cells by latrunculin A only achieved statistical significance in the presence of GSI (P=0.02) (Figure 1E).

K562 wild-type (WT) cells modified to express a BCMA–mCherry fusion protein, enabled simultaneous detection of BCMA and mCherry fluorescence. Treatment with GSI enhanced BCMA surface expression (Figure 2A). In an IncuCyte-based cytotoxicity assay, CAR T cells co-cultured with K562BCMA-mCherry cells, with or without GSI treatment, exhibited enhanced cytotoxicity compared to tumor-only and non-transduced T-cell co-cultures. This killing effect was more pronounced and sustained over time (Figure 2B, Suppl movie 1, Suppl movie 2). Upon GSI treatment, trogocytosis was significantly increased during co-culture with K562BCMA-mCherry cells, as evidenced by elevated BCMA MFI on CAR T cells at 30 min (P=0.007), 1 h (P<0.0001), 2 h (P<0.0001), and 6 h (P=0.0006) (Figure 2C). Confocal microscopy further confirmed the occurrence of trogocytosis (Figure 2D; Suppl movie 1). Notably, CD4+ CAR T cells exhibited significantly higher BCMA MFI at 10 min (P=0.01), 2 h (P=0.01), and 6 h (P=0.004) during co-culture with GSI-treated K562BCMA-mCherry cells than CD8+ CAR T cells (Figure 2E). To model fratricide, CAR T cells were co-cultured with K562BCMA-mCherry+GSI cells for 6 hours. CAR T cells that acquired BCMA via trogocytosis (hereafter referred to as CAR T Trogo+ cells) and those did not (CAR T cells) were sorted and subsequently re-challenged with naïve CAR T cells 24 hours (Figure 2F). The proportion of residual mCherry+ CAR T Trogo+ cells was significantly lower when co-cultured with naïve CAR T cells compared to the CAR T Trogo+ alone condition (P=0.002 for percentage, P=0.0005 for absolute counts; Figure 2G). Furthermore, the extent of fratricidal killing increased proportionally with higher CAR T: CAR T Trogo+ ratios (Figure 2H). Confocal imaging demonstrated fratricidal interactions between CAR T cells in the K562 BCMA-mCherry+GSI co-culture in time-lapse imaging (Figure 2I; Suppl movie 3).

**Figure 2A.**
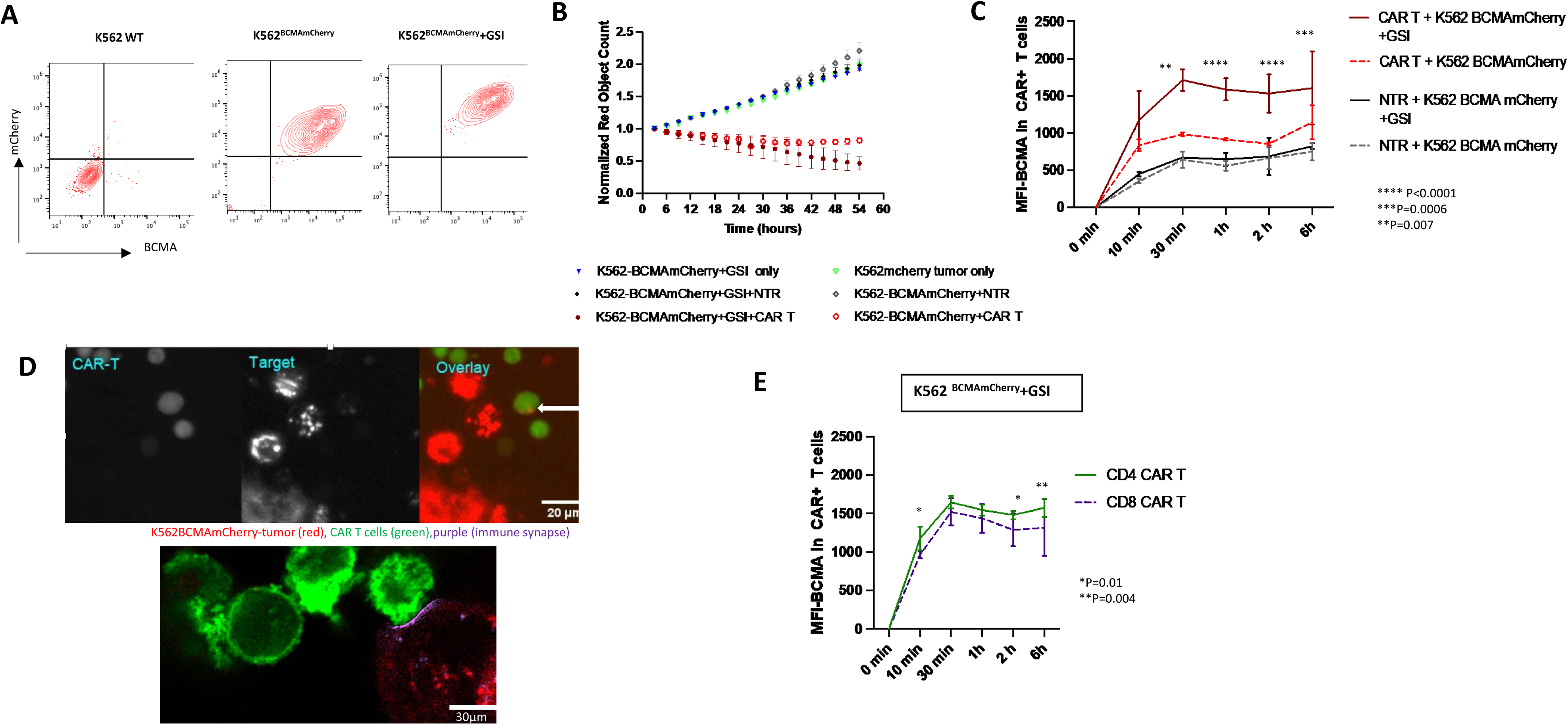
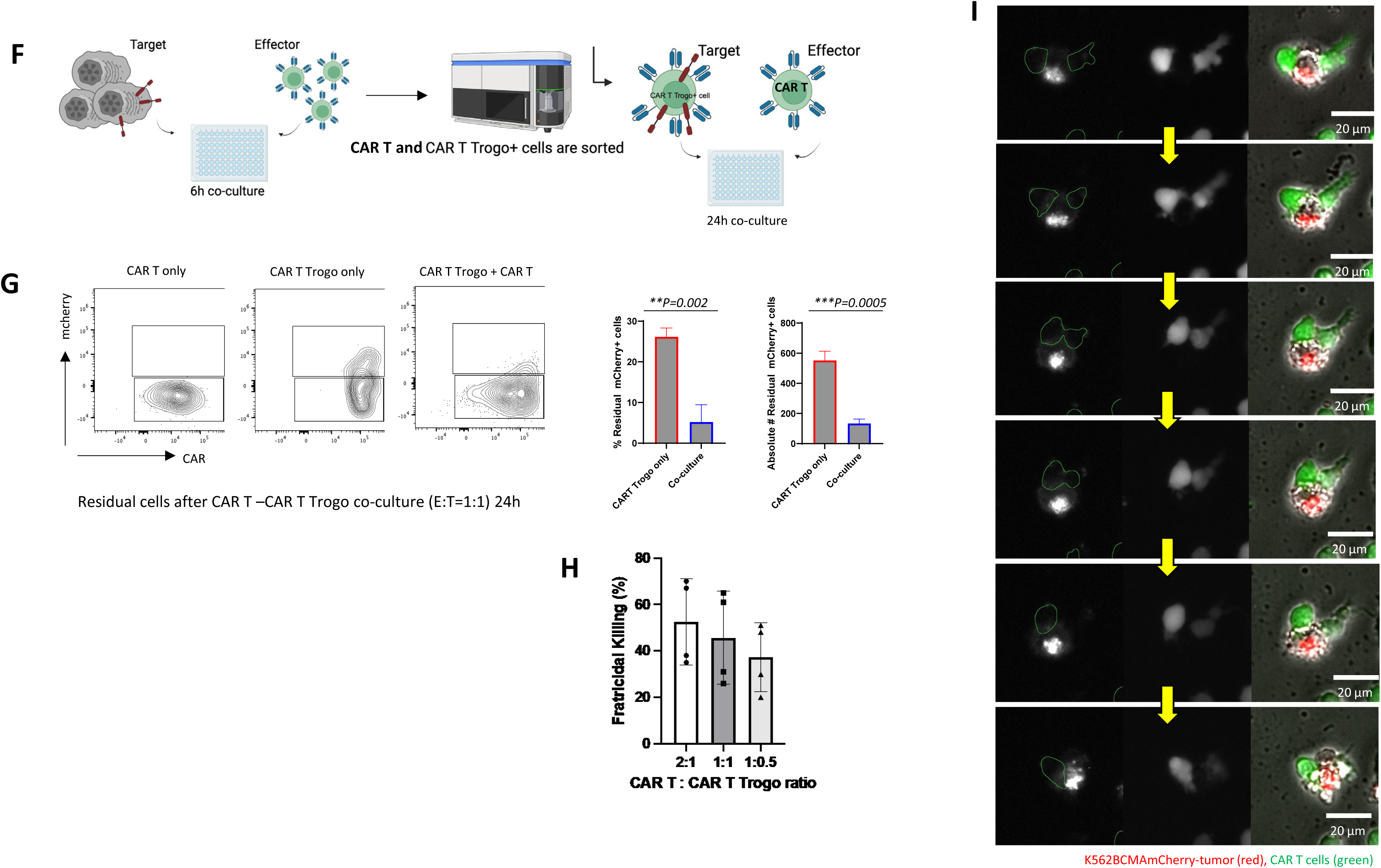
The BCMA and mCherry expression of K562BCMAmCherry +/−GSI **2B.** The normalized red object (mCherry) count in incucyte live cell analysis, images were taken every 3h after co-culture **2C.** The BCMA-MFI in CAR+ T and non-transduced T cells in co-cultures with K562 BCMAmCherry cell line +/−GSI (E:T=1:1) **2D.** Confocal images of trogocytosis (white arrow) after 2h of K562 BCMAmCherry+GSI and CAR T cells co-culture (E:T=1:1) (Top), the immune synapse is colored as purple (Bottom) **2E.** The trogocytosis in CD4 and CD8 CAR T cells with K562BCMAmCherry cell line + GSI (E:T=1:1). (E: Effector, GSI: Gamma secretase inhibitor, NTR: Non-transduced, T: Target) **Figure 2F**. The scheme of fratricide experiment. K562 BCMAmCherry + GSI co-cultured with CAR T cells for 6 hours (E:T=1:1) and CAR T Trogo cells were sorted and re-cultured with CAR T cells for 24 hours (E:T=1:1) **2G.** The flow plot showing mCherry expression after CAR T and CAR T Trogo cells (E:T=1:1) were co-cultured overnight, percentage and absolute count of residual mCherry+ cells in culture. **2H.** Percentage of fratricidal killing in different CAR T: CAR T Trogo ratios **2I.** The confocal images demonstrating fratricide in between CAR T cells in K562 BCMAmCherry+GSI and CAR T cells co-culture assay (E:T=1:1).The images were recorded in every 30 minutes. (E: Effector, GSI: Gamma secretase inhibitor, T: Target)

To assess the functional consequences of trogocytosis, sorted CAR T Trogo+ cells and CAR T cells were re-challenged with K562BCMA-mCherry+GSI cells and their cytotoxic activity was compared (Figure 3A). CAR T Trogo+ cells exhibited a reduced expansion profile during culture (Figure 3B; P=0.002). In a 24-hour cytotoxicity assay, both CAR T and CAR T Trogo+ cells demonstrated enhanced tumor cell killing (P<0.0001, respectively) however, the cytotoxic capacity of CAR T Trogo+ cells was significantly lower compared to CAR T cells (P=0.01) (Figure 3C). Despite this reduction in proliferative and cytolytic capacity, IFN-γ, TNF-α and IL-2 secretion remained comparable between the two groups following antigen re-challenge (Figure 3D), indicating that acute cytokine production was preserved. Phenotypic analysis revealed a significantly higher frequency of effector T (Teff) cells among both CD4+ and CD8+ CAR T Trogo+ T compared with their corresponding CAR T cell populations (P<0.0001, P=0.002). Furthermore, the T effector memory (Tem) subset was significantly enriched in CD4+ CAR T Trogo+ cells compared with CD4+ CAR T cells (P<0.0001) (Figure 3E). In CD4+ and CD8+ T cells, the frequency of exhaustion/activation markers, including PD-1+LAG-3+, PD-1+TIM-3+, and TOX+TIM-3+ co-expression, was significantly higher in CAR T Trogo+ cells compared to naïve CAR T cells after re-challange (P=0.007, P<0.0001, P=0.006 and P=0.004, P=0.03, P=0.006, respectively)(Figure 3F).

**Figure 3A.**
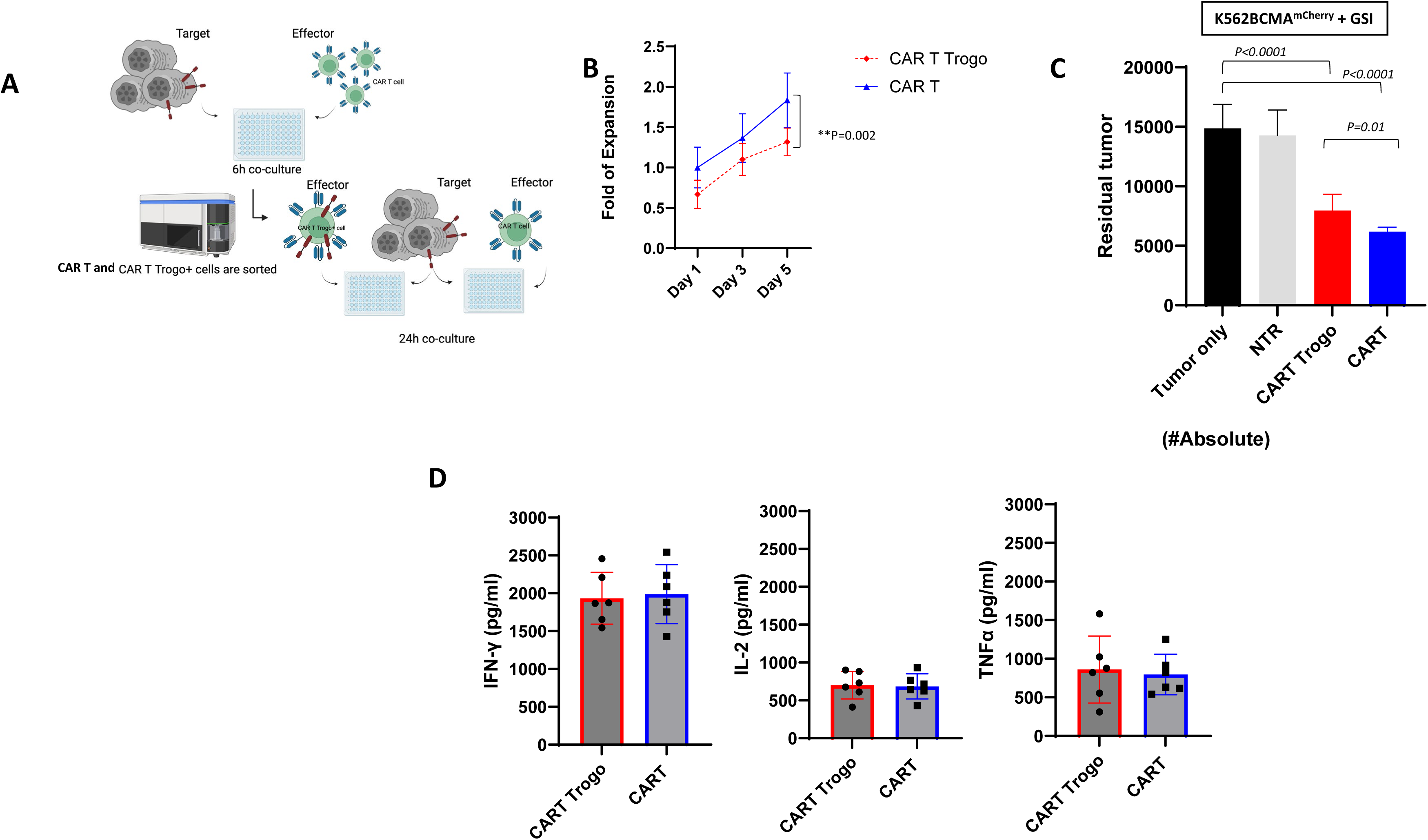
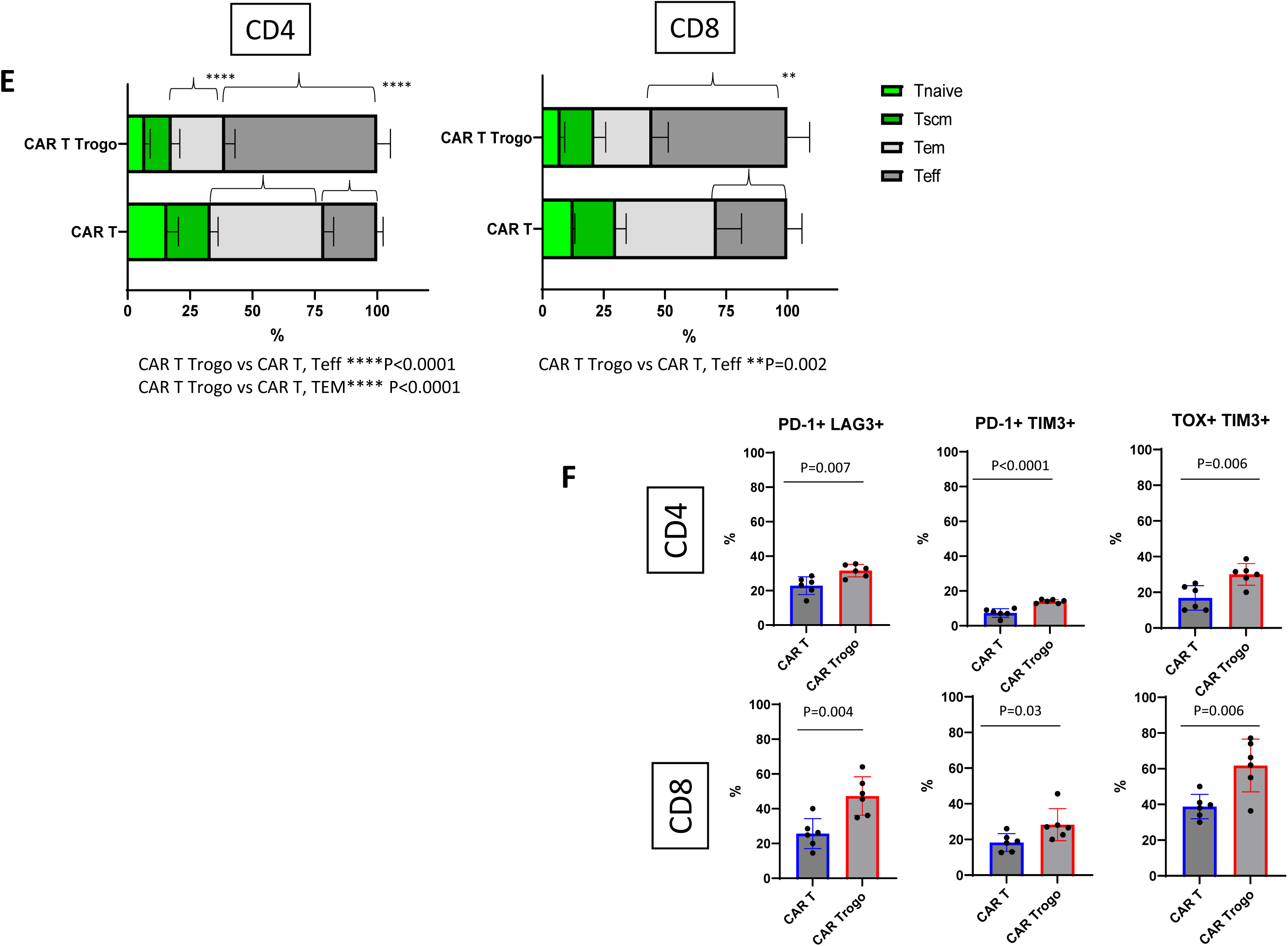
The scheme of tumor re-challenge experiment. K562 BCMAmCherry + GSI co-cultured with CAR T cells for 6 hours (E:T=1:1) and CAR T Trogo cells were sorted and re-challenged with K562 BCMAmCherry for 24h (E:T=1:1) **3B.** The fold of expansion of CAR T cells and CAR T Trogo cells after sorting (n=6) **3C.** Absolute Count of residual tumor cells after 24h tumor re-challenge (NTR: Non-transduced). **3D.** IFNγ, IL-2 and TNFα release after 24h tumor re-challenge. **3E.** Phenotype of all CD4 and CD8 CAR T *vs* CAR T Trogo cells following re-challenge. **3F.** Activation/Exhaustion markers (PD-1, LAG3, TIM-3, TOX) of CD4 and CD8 cells after tumor re-challenge

To define the molecular programs associated with trogocytosis, we performed integrated single-cell RNA sequencing (scRNA-seq) and paired T-cell receptor sequencing (scTCR-seq) on sorted CAR T Trogo+ and CAR T cells. Analysis of 2,880 high-quality T cells demonstrated partial transcription separation between two populations on UMAP, indicating that trogocytosis induces a distinct, yet overlapping, cellular state rather than generating a completely discrete population (Figure 4A). Differential expression analysis identified a broad transcriptional remodeling in CAR T Trogo+ cells, characterized by significant upregulation of genes associated with T-cell activation and effector function (*TNFRSF9, CRTAM, GZMB, LTA*), inflammatory cytokine production (*IL2, IL3, CSF1, CSF2*), chemokine signaling (*CCL1, CCL22, CXCL10*), cytoskeletal remodeling (*CDC42BPA, MYO1D, ARHGAP6*), and signaling pathways involved in cellular activation and differentiation (*RBPJ and TLR2*) (Figure 4B, D). Consistent with these findings, GSEA demonstrated significant enrichment of the hallmark TNFα signaling via NFκB pathway in CAR T Trogo cells (NES=2.37), indicating the activation of the TNFα/NF-κB inflammatory program represents a dominant transcriptional feature following trogocytosis (Figure 4C).

**Figure 4.**
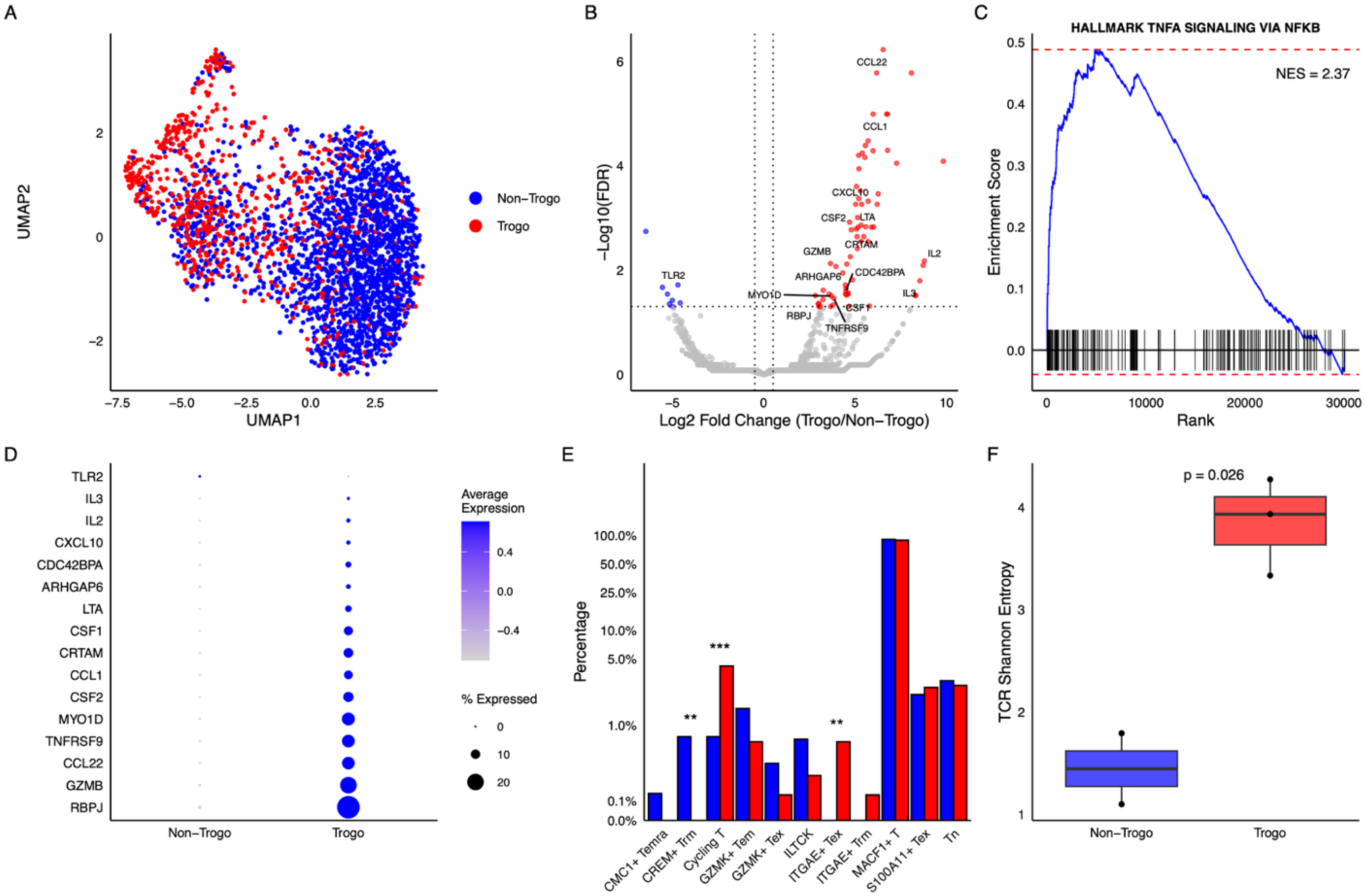
Integration of scRNA-seq and TCR-seq reveals the multi-omic signature of trogocytosis in CAR T cells. **A.** UMAP projection of 2,880 integrated T cells. The visualization is colored by FACS-sorted condition. **B.** Volcano plot comparing Trogo vs. Non-Trogo cells. Significance thresholds (dotted lines) are set at padj < 0.05 and |log2FC| > 0.5. Labeled genes represent a curated list of activation, exhaustion, and cytotoxicity markers. **C.** GSEA enrichment plot for the HALLMARK TNFA SIGNALING VIA NFKB pathway. The positive normalized enrichment score (NES = 2.37) confirms that NF-κB signaling is a primary transcriptomic hallmark of trogocytosis cells. **D.** Dot plot showing expression levels of significantly differentially expressed genes categorized by function: activation/effector (*TNFRSF9, CRTAM, GZMB, LTA*), cytokines and growth factors (*IL2, IL3, CSF1, CSF2*), chemokines (*CCL1, CCL22, CXCL10*), signaling and fate (*RBPJ, TLR2*), and cytoskeletal regulation and mechanics (*CDC42BPA, MYO1D, ARHGAP6*). Dot size represents the percentage of cells with detectable expression; color gradient indicates Z-scored average expression. **E.** Bar plot showing the proportion of scAtlasVAE-identified T cell subtypes on a pseudo-log percent scale. Asterisks indicate statistical significance via Fisher’s exact test (*p < 0.05, **p < 0.01, ***p < 0.001). **F.** Box plot of Shannon Entropy for the TCR repertoire. Each point represents an experimental sample. The significantly higher entropy in Trogo cells (p = 0.026) indicates a more diverse distribution of T-cell clonotypes.

To further investigate the phenoyptic composition of these cells, we projected the datasent onto the scAtlasVAE immune reference atlas. CAR T Trogo+ cells showed a significant enrichment of activated cytotoxic T-cell subsets, particularly Cyt(cycling) cells, together with increased representation of activated ITGAE-positive populations, whereas CAR T cells were relatively enriched for naïve/memory like subsets (Figure 4E). Integration of paired TCR sequencing further revealed significantly greater TCR repertoire diversity in CAR T Trogo+, as demonstrated by increased Shanon entropy (P=0.026) indicating that trogocytosis is associated with a broader distribution of T-cell clonotypes rather than expansion of a limited number of dominant clones (Figure 4F).

Finally, we analyzed data from two Phase I clinical trials evaluating FCARH143 CAR T cells administered without γ-secretase inhibitor (GSI; NCT03338972) and with GSI (NCT03502577) (Figure 5A). The normalized mean fluorescence intensity (MFI) of BCMA on CAR T cells for individual patients did not significantly differ between peripheral blood (PB) and bone marrow (BM) samples on days 14 or 28 in either trial (Figure 5B). CD4⁺ CAR T cells exhibited a trend toward higher BCMA trogocytosis at all evaluated time points (PB/BM days 14 and 28); however, these differences were not statistically significant (Figure 5C). Soluble BCMA (sBCMA) levels at days 14 and 28 were comparable between the two trials (Figures 5D). Patients with shorter CAR T cell persistence tended to have higher normalized BCMA MFI on their CAR T cells, although this trend did not reach statistical significance (Figure 5E). Notably, the extent of trogocytosis did not correlate with duration of response in either study (Figure 5F).

**Figure 5A.**
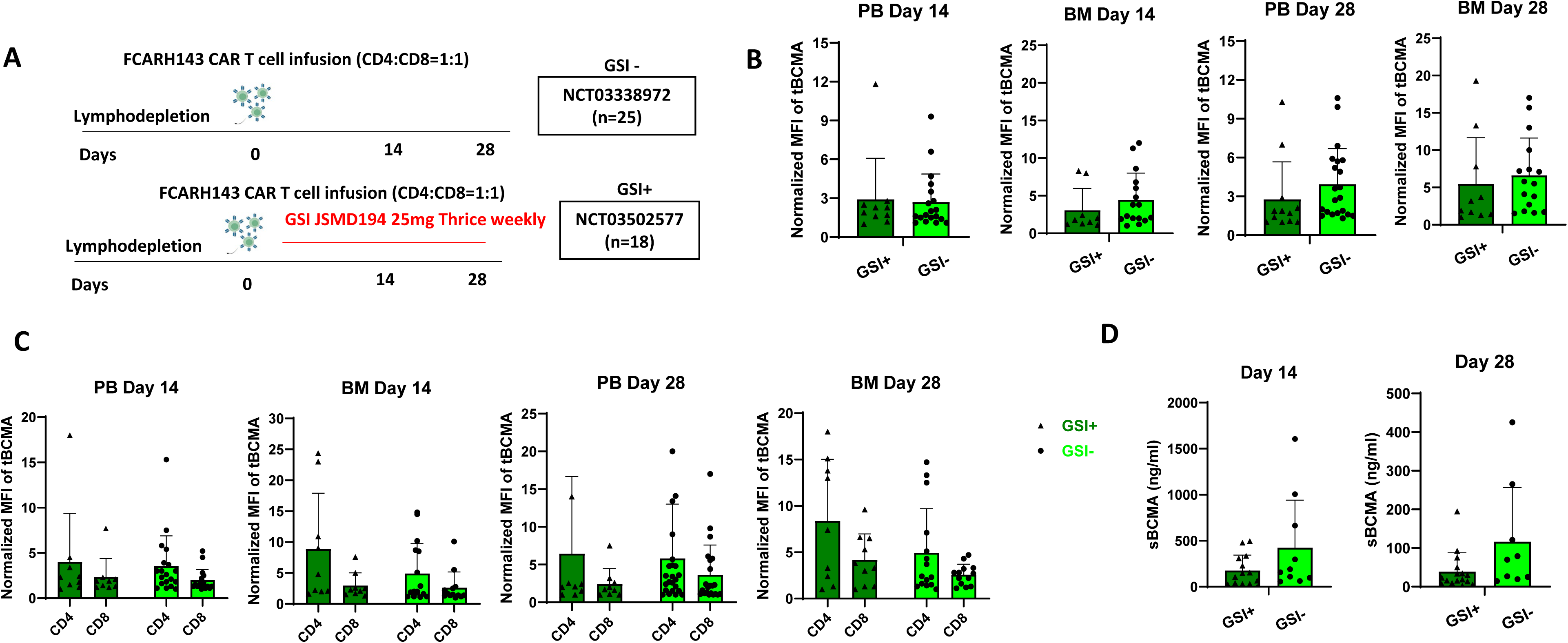
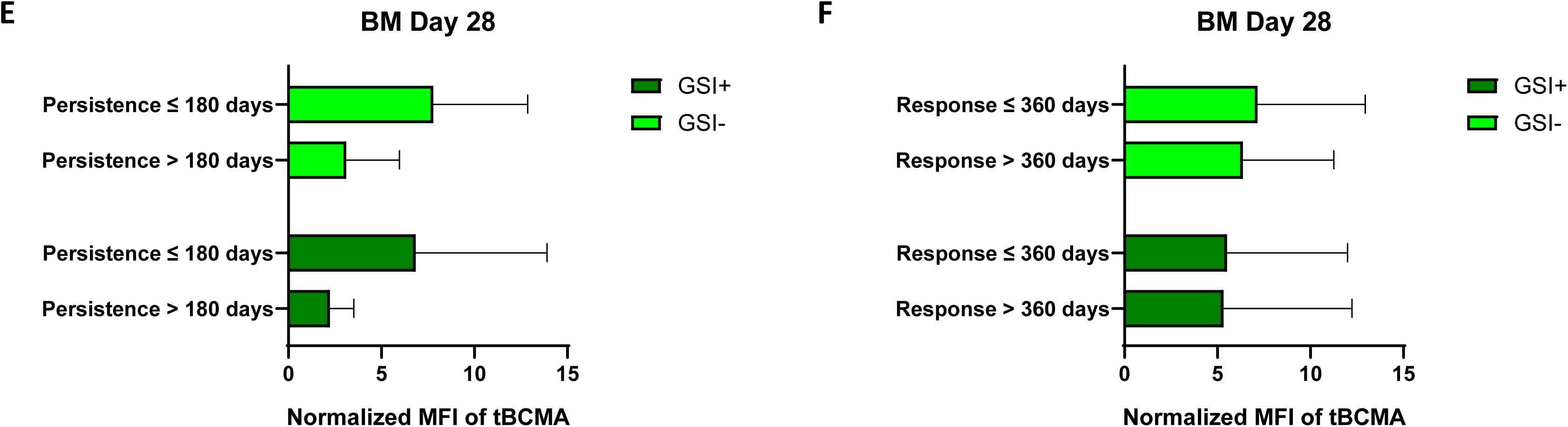
The clinical design of two Phase I trials (NCT03338972-GSI negative; NCT03502577-GSI positive) **5B.** Normalized geometric mean fluorescent intensity (MFI) of trogocytosed BCMA (tBCMA) on CAR T cells in peripheral blood (PB) and bone marrow (BM) samples from day 14 and day 28 in GSI negative and GSI positive patient groups (PB Day 14 GSI+ (n=10) vs GSI-(n=14); BM Day 14 GSI+ (n=9) vs (n=16); PB Day 28 GSI+ (n=12) vs GSI-(n=21); BM Day 28 (n=10) vs GSI-(n=16) **5C.** Normalized geometric mean fluorescent intensity (MFI) of trogocytosed BCMA (tBCMA) on CD4 and CD8 CAR T cells in peripheral **Figure 5E**. The comparison of CAR persistence and normalized MFI of tBCMA in between GSI+ and GSI-patient groups; short persistence is of CAR with qPCR≤180 days **5F.** The comparison of duration of response and normalized MFI of tBCMA in between GSI+ and GSI-patient groups; short duration of response is ≤360 days

## Discussion

In particular, trogocytosis is characterized by ‘nibbling’ on the part of the cell membrane of the donor cell rather than whole cell-phagocytosis and can occur between multiple cell types (20,29). CAR T cells can acquire target antigens from tumor cells and display them on their surface (17,30). Pharmacologic inhibition of BCMA cleavage and the ensuing increase in antigen density enhances BCMA trogocytosis by anti-BCMA CAR T cells, results in increased antigen transfer, CAR molecule exchange, fratricide and progressive functional impairment of CAR T cells. This finding reveals that trogocytosis represents a dynamic biological process that reshapes CAR T cell phenotype beyond simple antigen acquisition. CAR T cells acquiring BCMA exhibit reduced proliferative capacity, diminished cytotoxicity upon re-challenge, increased expression of activation and exhaustion markers, enrichment of inflammatory TNFα/NF-κB transcriptional programs, and greater TCR repertoire diversity. Together, these findings identify an unanticipated consequence of GSI-mediated BCMA enhancement where increased antigen density drives trogocytosis which in turn contributes to CAR T cell dysfunction.

The successful blockage of BCMA cleavage by γ secretase inhibitors (GSI)s has emerged as an attractive strategy to improve the efficacy of BCMA-directed immunotherapy by uniformly increasing target antigen density. Our previous preclinical study resulted in enhanced CAR T cell efficacy (15) and supported a Phase I clinical trial. This first-in-human study of a BCMA CAR T cell therapy in combination with oral GSI demonstrated improved CAR T cell recognition and anti-myeloma activity (16), however, the current findings reveal an important trade-off. The same increase in antigen density substantially augmented BCMA transfer to CAR T cells in time dependent manner, suggesting that higher antigen availability increases opportunities for immune synapse mediated membrane exchange.

The mechanism of tumor antigen transfer to CAR immune cells via trogocytosis is not well understood but is thought to be related to the synaptic formation, antigen density, and affinity or CAR molecules with the target antigens (31). Trogocytosis is not specific to certain antigens but is intrinsically linked to the target recognized by the CAR. Similar to our study, CD19-directed CAR T cells mediated the transfer of CD19, while mesothelin (MSLN)-targeting CAR T cells facilitated the transfer of MSLN (17,32). Latrunculin A blocked trogocytosis confirming that immune synapse formation is necessary between BCMA CAR T and target (17). Our findings further extend previous observations by demonstrating that target antigen density is a major determinant of the magnitude of trogocytosis. Across MM cell lines with varying baseline BCMA expression, GSI-mediated upregulation of surface BCMA consistently resulted in greater antigen acquisition by CAR T cells, indicating that increased antigen availability directly promotes membrane transfer during CAR-target engagement. While analysis of samples from our two Phase I clinical trials did not confirm the impact of GSI on target trogocytosis, the studies were not designed to evaluate trogocytosis. The earliest available time point for sample collection was day 14 post-infusion. Given that trogocytosis is a dynamic and transient process in vivo (20), it is likely that key events occurred at earlier time points. Moreover, by day 14, disease burden in most patients was already substantially reduced due to CAR T cell activity, potentially confounding the detection and interpretation of trogocytosis-related phenomena. Isolating the impact of trogocytosis is confounded in the trials, because clinical response is a multifactorial outcome influenced by various patient– and therapy-related parameters. Identifying the impact of trogocytosis in clinical settings requires a prospective trial design to facilitate a comprehensive multivariable analysis.

Current research has primarily focused on the phenomenon of antigen transfer, whereby target antigens are acquired by CAR T cells, rendering them susceptible to recognition and elimination by other CAR T cells—commonly referred to as ‘fratricide.’ This process may have important implications for CAR T cell persistence and therapeutic durability. Using a K562BCMA-mCherry model, we confirmed that GSI treatment amplified CAR T cell-mediated cytotoxicity and trogocytosis, with direct visualization of antigen acquisition and fratricide events via confocal and time-lapse imaging. Furthermore, CAR T Trogo+ cells displayed reduced proliferative expansion and impaired cytotoxicity during tumor re-challenge despite maintaining comparable cytokine secretion of IFN-γ, TNF-α, and IL-2, accompanied by an increased expression of activation/exhaustion markers. This functional dissociation suggests that trogocytosis does not abolish antigen responsiveness but instead shifts CAR T cells toward a chronically activated state in which inflammatory signaling in preserved despite impaired proliferative and cytolytic capacity. Consistent with this interpretation, CAR T Trogo+ cells displayed increased expression of PD-1, TIM-3, LAG-3, and TOX, enrichment of effector/effector memory phenotypes, and activation of TNFα/NF-κB inflammatory transcriptional programs by scRNA-seq. In a study of CD19 CAR NK cells, trogocytosis of CD19 was shown to enhance early activation, evidenced by increased IFN-γ production and CD107a expression. However, this initial activation was followed by CAR-NK hyporesponsiveness, impaired long-term antitumor activity, and metabolic dysfunction associated with reduced glycolytic fitness (18). Importantly, consistent with Zhai et al.,we found that tumor cells are also able to acquire CAR molecules and ‘express’ them on the cell surface, the transfer leads to the loss of the CAR molecules in CAR T cells, impaired activity and antigen masking, forming drug resistance against CAR T cells (30).

Single-cell transcriptomic analysis provided mechanistic insight into the biological state associated with trogocytosis. Rather than forming a completely distinct population, Trogo+ CAR T cells occupied a transcriptionally shifted state characterized by activation of inflammatory, cytotoxic, and cytoskeletal remodeling programs. Among the enriched pathways, TNFα signaling via NF-κB emerged as the dominant transcriptional signature, consistent with sustained antigen receptor signaling and activation-induced stress. Upregulation of genes including *TNFRSF9, CRTAM, GZMB, CXCL10, CSF2*, and *CDC42BPA* further supports a phenotype of heightened activation accompanied by extensive immune synapse remodeling. Interestingly, projection onto the scAtlasVAE reference atlas demonstrated enrichment of activated cytotoxic and cycling T-cell subsets, whereas paired TCR sequencing revealed greater repertoire diversity rather than oligoclonal expansion. These findings suggest that trogocytosis is not restricted to a limited number of highly expanded clones but instead represents a broadly distributed adaptive response affecting multiple CAR T-cell clonotypes.

The underlying mechanisms driving trogocytosis and strategies to effectively moderate it has not been fully characterized. In recent studies, it was revealed that cholesterol-25-hyroxylase (CH25H) is a negative regulator of cell membrane cholesterol and plays an important role in membrane stability and fluidity (33). Tumor-derived factors (TDFs), such as PGE2 and VEGF, down-regulate the expression of CH25H by activating the transcription factor 3 (ATF3), promoting trogocytosis between CAR-T cells and tumor cells, while also limiting the viability and anti-tumor activity of CAR-T cells (34). Dietze et al proposed cysteine protease cathepsin B (CTSB) as a driver of CAR-mediated trogocytosis (35). Now several approaches have been proposed to overcome trogocytosis such as low-affinity CARs, dual CAR systems, inhibitor CAR for self-recognition, down-regulation of ATF3 and up-regulation of CH25H and dynamic regulation of CAR molecular expression (25). Interestingly, we report that CD4 CAR T cells exhibited higher levels of trogocytosis of the target antigen compared to CD8 CAR T cells in our study. The observed trend toward higher levels of BCMA trogocytosis in CD4⁺ CAR T cells compared to CD8⁺ CAR T cells warrant further investigation to elucidate underlying mechanisms and potential clinical implications. Previous studies have demonstrated that trogocytosis in CD4^+^ T cells sustains intracellular signaling and promotes prolonged T-cell activation following antigen engagement, suggesting that intrinsic differences in immune synapse dynamics or signaling may render CD4^+^ CAR T cells more susceptible to antigen acquisiton (36, 37) Furthermore, CD4⁺ helper T cells play a critical role in supporting CAR T-cell expansion, persistence, and the orchestration of antitumor immune responses. Preferential trogocytosis within this compartment may therefore disproportionately contribute to chronic antigen stimulation, activation-induced dysfunction, and progressive loss of CAR T-cell fitness. This concept is supported by our findings demonstrating increased effector differentiation, higher expression of activation/exhaustion markers, and inflammatory transcriptional reprogramming in CAR T Trogo⁺ cells.

Collectively, our findings reveal that enhanced antigen density is a double-edged sword. Enhanced CAR T cell activation can more effectively eliminate target tumor cells, but simultaneously amplifies trogocytosis-mediated antigen acquisition, fratricide, inflammatory transcriptional reprogramming and progressive functional impairment of CAR T cells. These results highlight the need for strategies that preserve the benefits of increased antigen expression while limiting trogocytosis-mediated dysfunction. More broadly, our data highlight the importance of comprehensively characterizing any therapeutic intervention that modulates target antigen expression to define its optimal timing, dose, and duration when administered in combination with adoptive cellular immunotherapies.

Although our Phase I clinical trials were not specifically designed to evaluate the relationship between GSI administration and trogocytosis, we were able to demonstrate that BCMA trogocytosis occurs in patients receiving BCMA CAR T-cell therapy. Notably, these findings may also have implications beyond CAR T-cell therapy. For example, recent clinical studies combining GSI with BCMA-directed bispecific antibodies have not demonstrated the anticipated improvement in clinical outcomes (38), raising the possibility that enhanced trogocytosis resulting from increased antigen density may partially attenuate the therapeutic benefit of this combination. While this hypothesis requires prospective validation, it underscores the need to evaluate trogocytosis as a potential biological consequence whenever antigen-enhancing strategies are incorporated into immune-based therapies. Future prospective clinical studies incorporating optimized longitudinal sampling can be pursued to define the kinetics of trogocytosis, determine its impact on treatment outcomes, and establish the optimal timing, duration, and frequency of GSI administration to maximize the efficacy and durability of BCMA CAR T-cell therapy.

## Acknowledgements

This research was supported by the National Institutes of Health, National Cancer Institute grant P01 CA018029; The Leukemia and Lymphoma Society Specialized Center of Research program. Research reported in this publication was also supported by the NCI/NIH Award P30CA240139. We wish to acknowledge Flow Cytometry (FSCR) Shared Resources at Sylvester Comprehensive Cancer Center; We thank Cellular Imaging Shared Resource-Hoku West-Foyle and Lena Schroeder-RRID:SCR_022609 of the Fred Hutch/University of Washington/Seattle Children’s Cancer Consortium (P30 CA015704). Figure 2A and figure 3A are created in BioRender.

