## Supplementary figures for "Enhanced BCMA Antigen Density Increases Trogocytosis and Attenuates CAR T cell Function"

### Slide 1
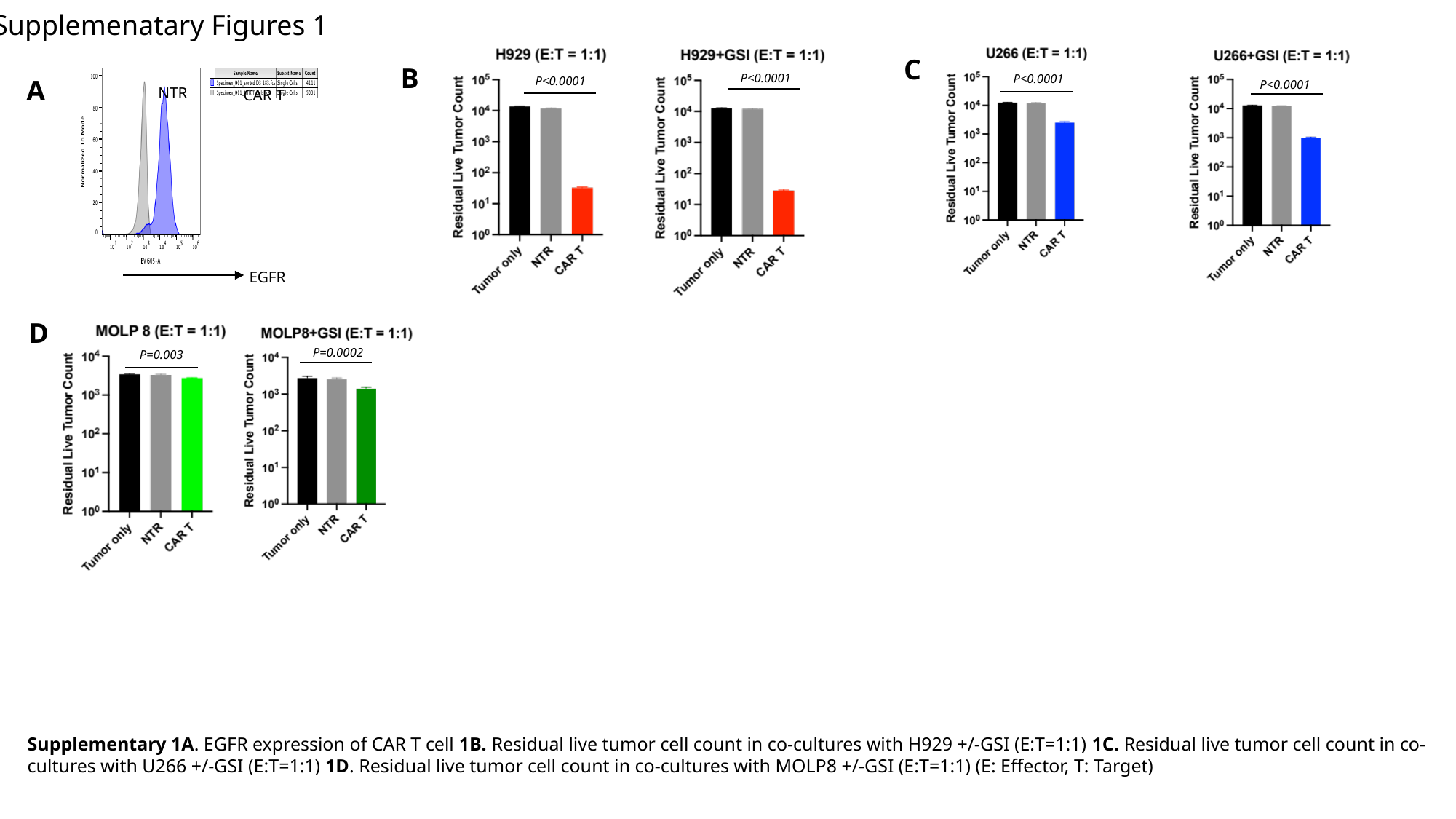

Supplemenatary Figures 1
P<0.0001
P<0.0001
C
P=0.003
P=0.0002
B
P<0.0001
A
NTR
CAR T
EGFR
P<0.0001
D
Supplementary 1A. EGFR expression of CAR T cell 1B. Residual live tumor cell count in co-cultures with H929 +/-GSI (E:T=1:1) 1C. Residual live tumor cell count in co-cultures with U266 +/-GSI (E:T=1:1) 1D. Residual live tumor cell count in co-cultures with MOLP8 +/-GSI (E:T=1:1) (E: Effector, T: Target)

### Slide 2
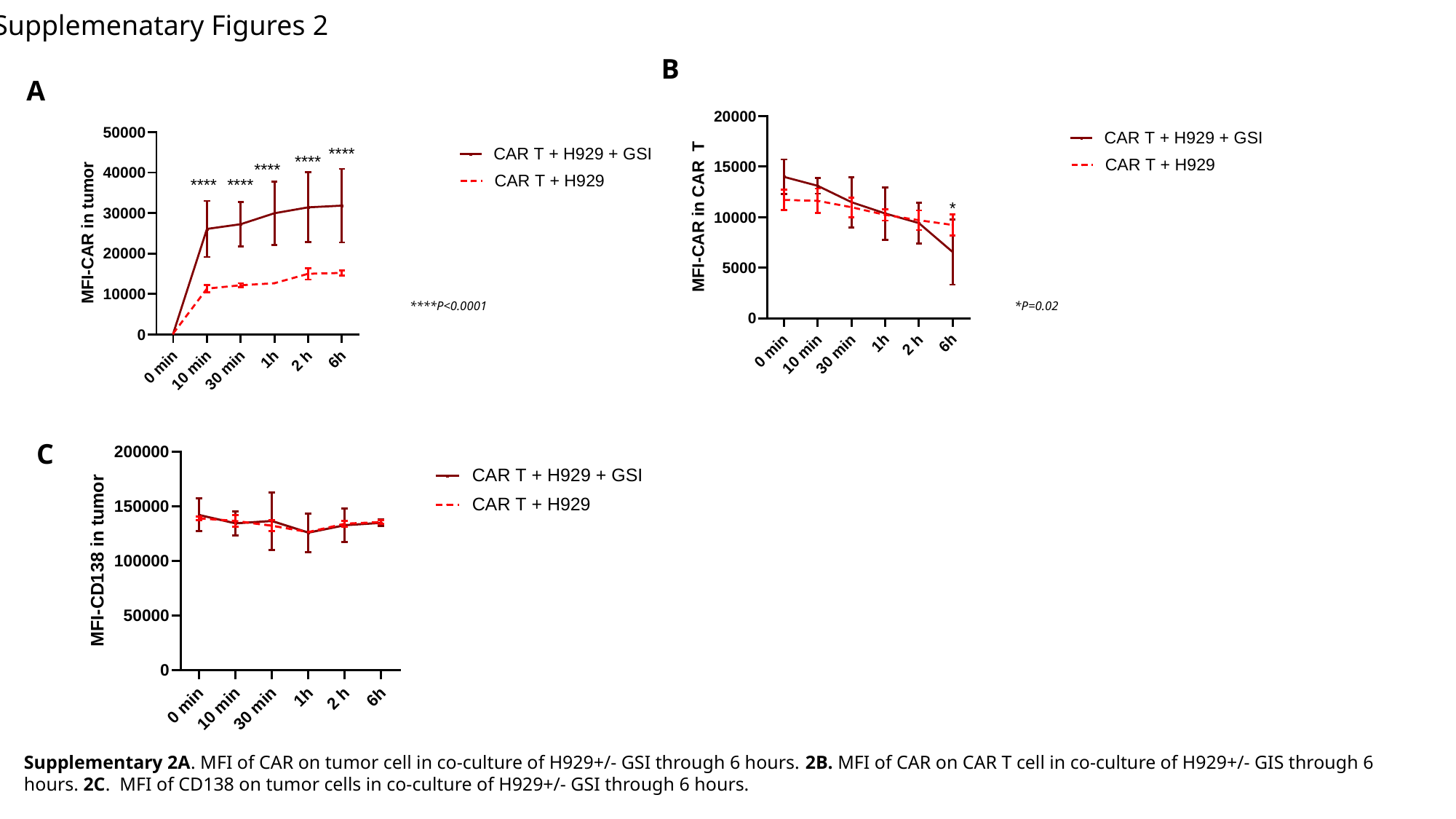

Supplemenatary Figures 2
B
A
****P<0.0001
*P=0.02
C
Supplementary 2A. MFI of CAR on tumor cell in co-culture of H929+/- GSI through 6 hours. 2B. MFI of CAR on CAR T cell in co-culture of H929+/- GIS through 6 hours. 2C. MFI of CD138 on tumor cells in co-culture of H929+/- GSI through 6 hours.

### Slide 3
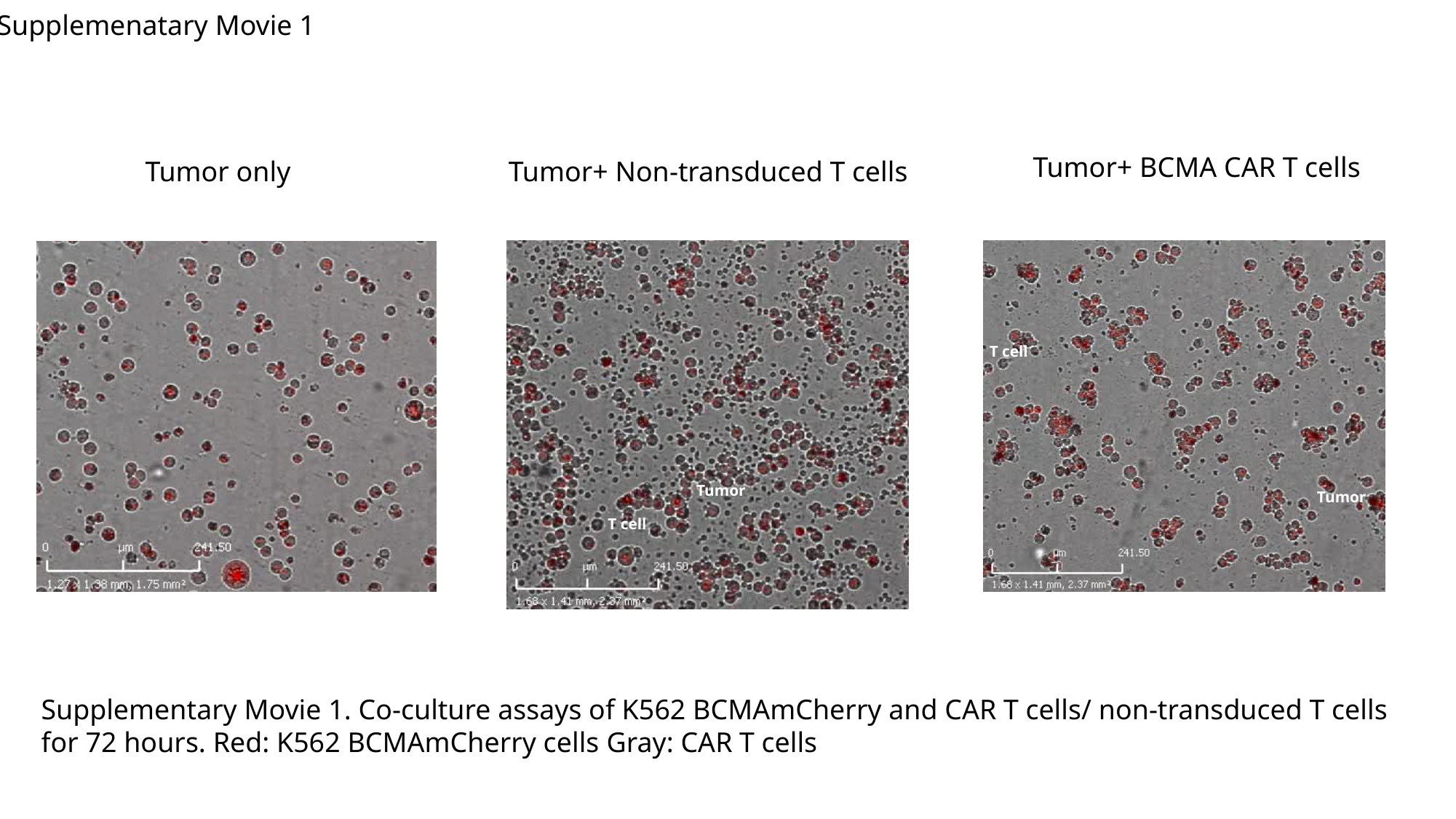

Supplemenatary Movie 1
Tumor+ BCMA CAR T cells
Tumor only
Tumor+ Non-transduced T cells
T cell
Tumor
Tumor
T cell
Supplementary Movie 1. Co-culture assays of K562 BCMAmCherry and CAR T cells/ non-transduced T cells for 72 hours. Red: K562 BCMAmCherry cells Gray: CAR T cells

### Slide 4
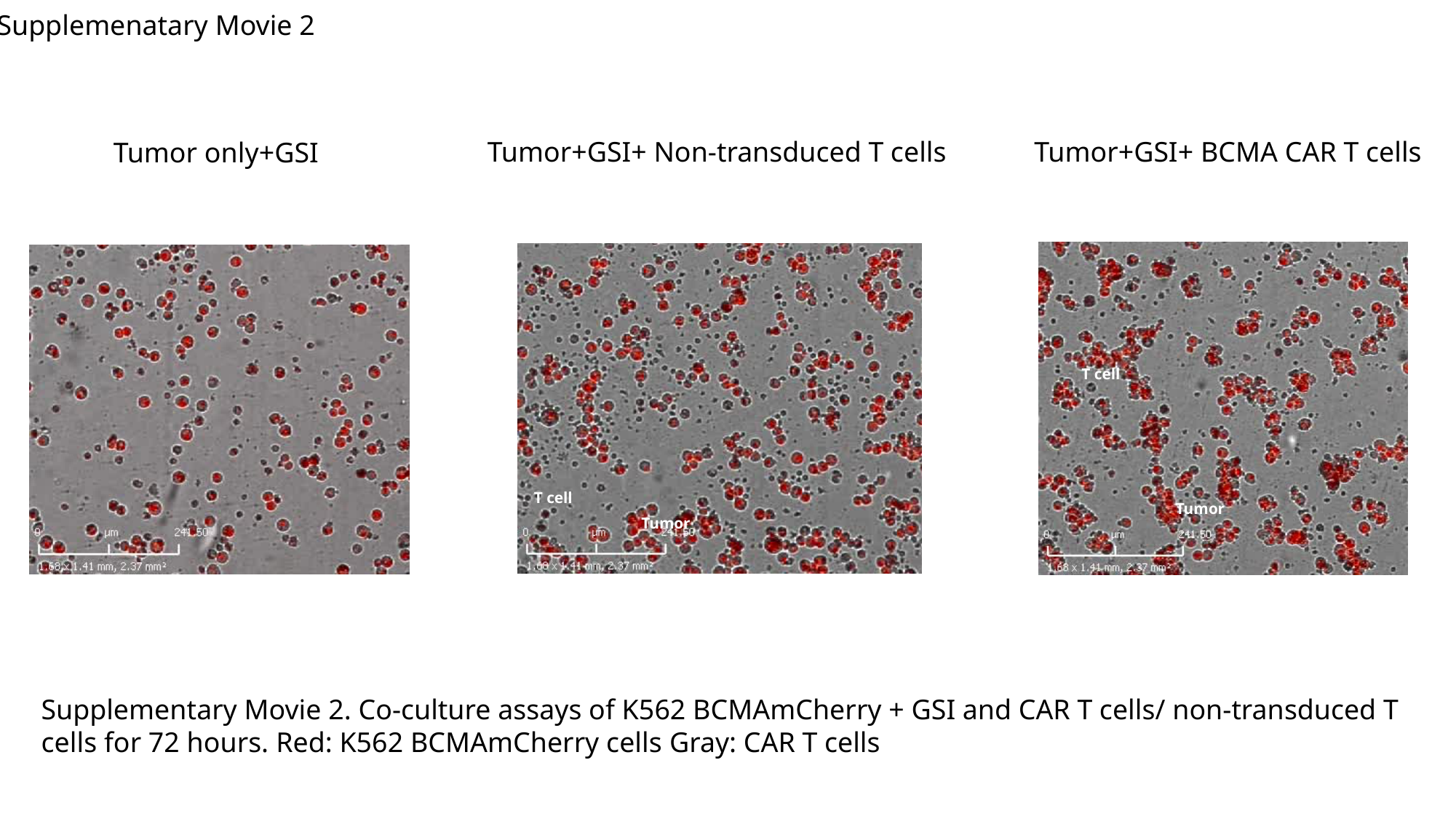

Supplemenatary Movie 2
Tumor+GSI+ Non-transduced T cells
Tumor+GSI+ BCMA CAR T cells
Tumor only+GSI
T cell
T cell
Tumor
Tumor
Supplementary Movie 2. Co-culture assays of K562 BCMAmCherry + GSI and CAR T cells/ non-transduced T cells for 72 hours. Red: K562 BCMAmCherry cells Gray: CAR T cells

### Slide 5
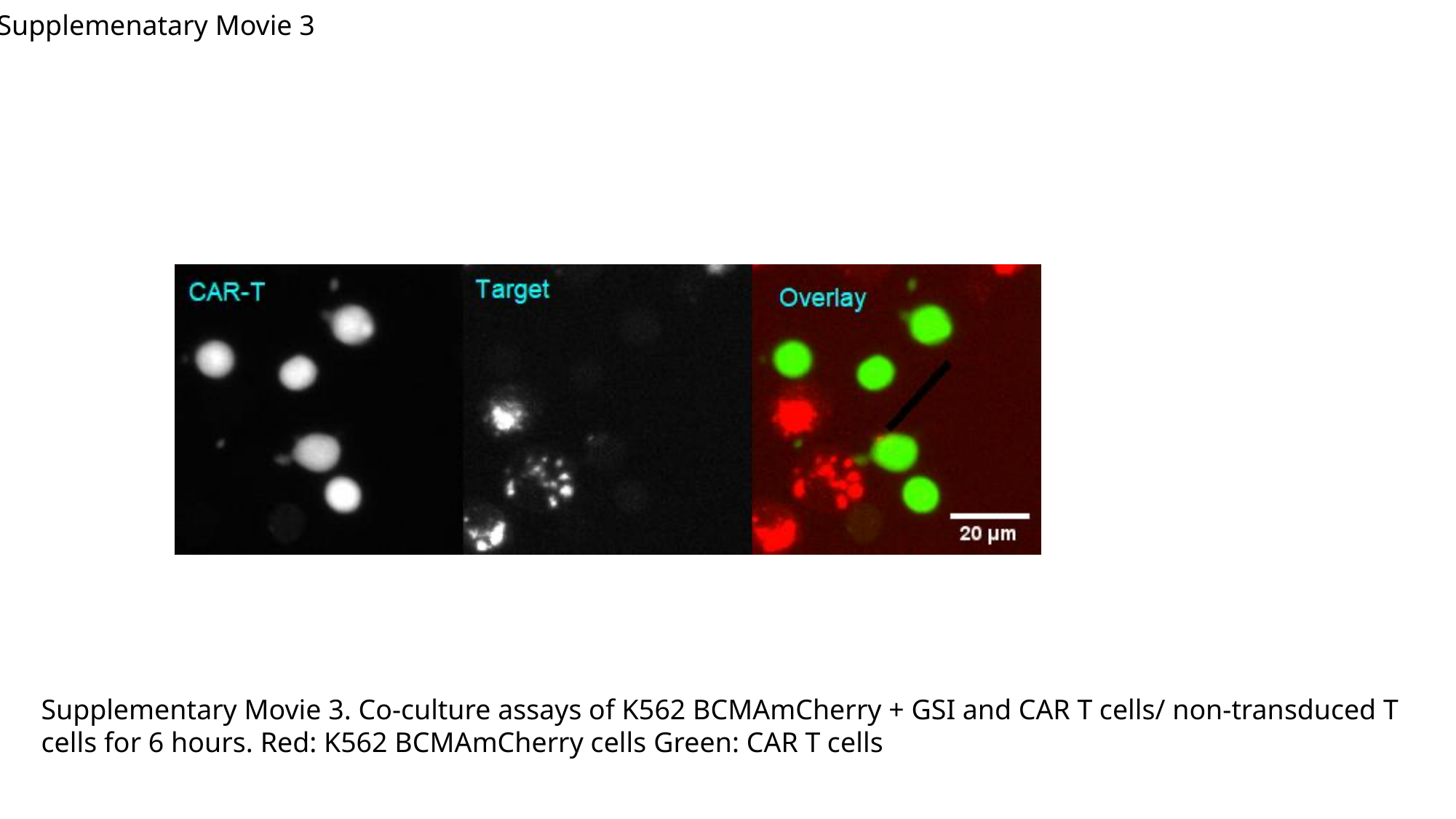

Supplemenatary Movie 3
Supplementary Movie 3. Co-culture assays of K562 BCMAmCherry + GSI and CAR T cells/ non-transduced T cells for 6 hours. Red: K562 BCMAmCherry cells Green: CAR T cells

### Slide 6
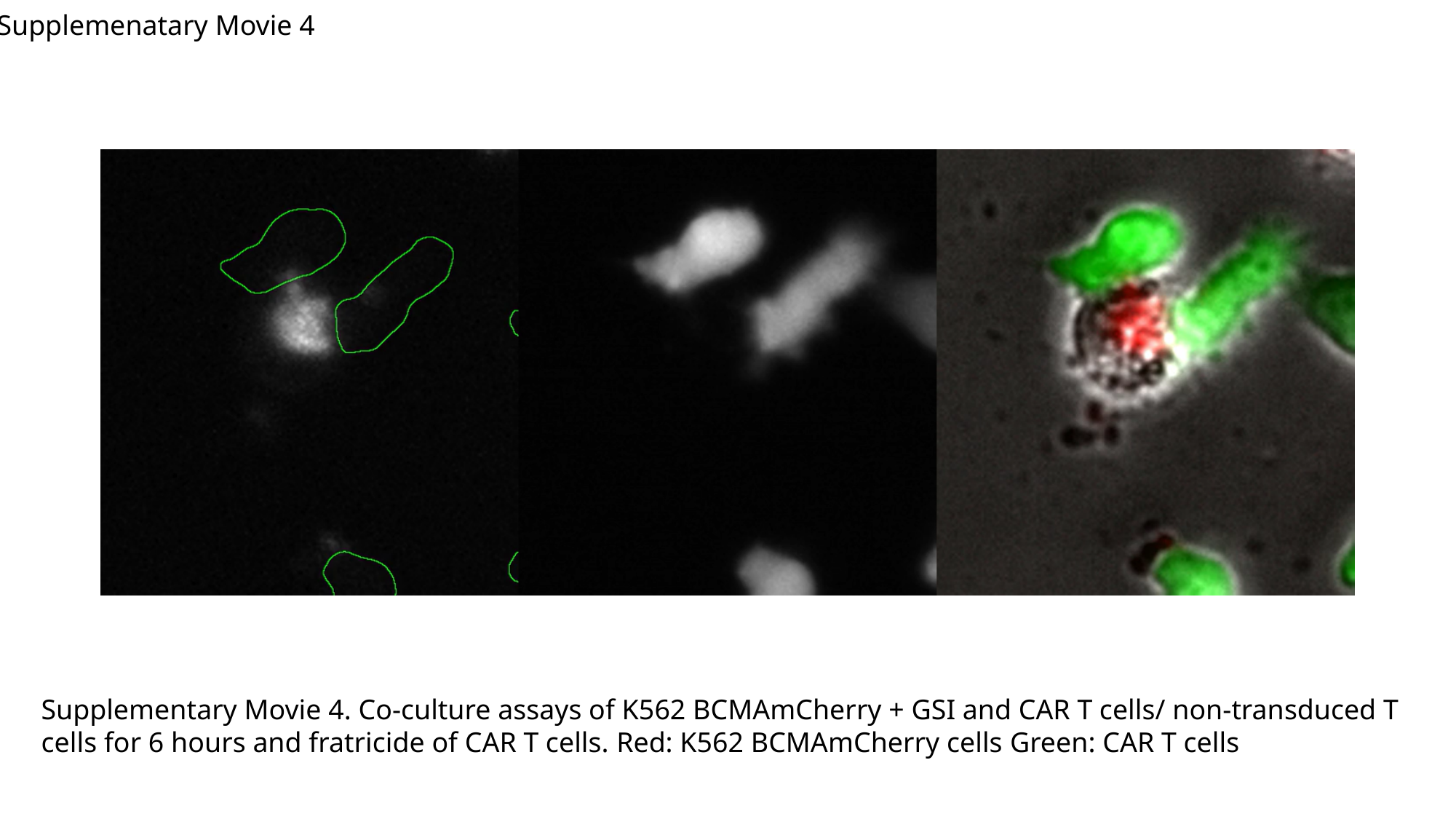

Supplemenatary Movie 4
Supplementary Movie 4. Co-culture assays of K562 BCMAmCherry + GSI and CAR T cells/ non-transduced T cells for 6 hours and fratricide of CAR T cells. Red: K562 BCMAmCherry cells Green: CAR T cells
